# Oncostatin M orchestrates collective epithelial migration via HIF1A activation

**DOI:** 10.1101/2025.09.26.678830

**Authors:** Ian C. McLean, Sean M. Gross, Tiera A. Liby, Mark A. Dane, Daniel S. Derrick, Laura M. Heiser

## Abstract

Extracellular signals strongly influence cell behavior, yet the mechanisms by which specific ligands mediate changes in phenotype remain unclear. The cytokine Oncostatin M (OSM) regulates homeostasis, wound healing, inflammation, and cancer progression. We previously found that OSM induces collective cell migration (CCM), a process where cells move as cohesive units while retaining cell-cell contacts, in MCF10A mammary epithelial cells. Here, we investigated how OSM drives CCM by comparing its effects with those elicited by epidermal growth factor (EGF) and interferon gamma (IFNG), defining ligand-specific phenotypes and molecular networks. Integrative transcriptomic and proteomic analyses identified hypoxia-inducible factor-1 (HIF1A) and signal transducer and activator of transcription 3 (STAT3) as central regulators of OSM responses. Functional validation revealed that HIF1A drives transcriptional programs associated with hypoxia, metabolic reprogramming, and immune pathways. Complement signaling emerged as a downstream effector of HIF1A, and its inhibition disrupted OSM-induced clustering and CCM. These findings establish a mechanistic link between OSM signaling, HIF1A activation, and CCM, demonstrating how cytokine-driven transcriptional reprogramming coordinates epithelial migration. Analysis of public breast cancer data suggests this pathway is active in human tumors and may contribute to tissue remodeling, repair, and metastasis.

## Introduction

Collective cell migration (CCM) is a highly coordinated process in which groups of cells migrate together, typically while maintaining cell-cell junctions and collective polarity.^1^ Unlike single-cell migration, which occurs independently, CCM depends on sustained intercellular communication and mechanical coupling between cells to maintain collective integrity and directionality.^2^ This process enables cohesive cellular groups to collectively respond to environmental cues, facilitating complex behaviors essential for tissue morphogenesis, wound healing, and immune responses.^1^ In developmental contexts, CCM plays a pivotal role in shaping organ architecture. For example, during the formation of neuronal streams in the developing brain, chains of neuroblasts migrate collectively from the subventricular zone to the olfactory bulb.^3^ Similarly, during mammary gland development, epithelial cells undergo branching morphogenesis, a process driven by collective migration that ensures the proper organization of ducts and alveoli.^4^ In wound healing, keratinocytes at the wound margin migrate collectively, closing the wound while preserving the integrity of the epithelial sheet.^5^ Despite its central role in diverse biological processes, the molecular mechanisms that coordinate CCM remain incompletely understood.

CCM involves diverse signaling networks and cytoskeletal dynamics,^2^ including maintenance of adherens junctions and coordinated cytoskeletal remodeling.^6^ Cell-cell adhesion molecules, such as E-cadherin, ensure physical cohesion among migrating cells, while integrins mediate interactions with the extracellular matrix, providing traction and directional guidance.^7^ Rho family GTPases regulate cytoskeletal rearrangements, enabling cells to generate the mechanical forces required for migration.^8^ Furthermore, growth factors and cytokines in the microenvironment, such as transforming growth factor-beta (TGF-β) and epidermal growth factor (EGF), have been shown to provide spatial and temporal cues that guide CCM.^9,10^ These signaling molecules activate downstream pathways, including MAPK, PI3K-AKT, and JAK-STAT, to modulate cellular polarity, adhesion, and motility.^11,12^

Beyond its roles in normal physiology, CCM has been implicated in cancer progression, particularly in metastasis. In breast cancer, tumor cells can adopt a collective migratory mode to invade surrounding tissues and disseminate to distant sites.^13^ The maintenance of partial junctional integrity during CCM facilitates coordinated movement through the stroma,^14,15^ while clusters of cancer cells exhibit improved survival during intravasation and extravasation, potentially due to retention of intercellular signaling and adhesion molecules that protect against anoikis.^16^ Additionally, these clusters may carry supportive stromal or immune cells that facilitate colonization at distant sites.^17–19^ Collective migration in cancer is often associated with epithelial-to-mesenchymal transition (EMT), characterized by downregulation of epithelial markers (e.g., E-cadherin) and upregulation of mesenchymal markers (e.g., vimentin).^20^ Environmental cues such as interleukin-6 (IL-6) and hypoxia further promote collective invasion by enhancing tumor cell motility and stromal remodeling.^21,22^

Among the diverse extracellular signals that regulate collective cell migration, Oncostatin M (OSM), an IL-6 family member, has recently been linked to the induction of CCM in epithelial cells.^23^ OSM signals through a heterodimeric receptor composed of OSMRβ and gp130, activating downstream pathways including JAK-STAT, MAPK, and PI3K-AKT.^24,25^ In mammary epithelial cells, OSM promotes phenotypic characteristics associated with collective migration, including enhanced motility, junctional remodeling, and the formation of cohesive cellular clusters.^23^ These observations suggest that OSM may act as a microenvironmental cue that drives CCM, making OSM-treated epithelial cells a valuable model for dissecting the molecular mechanisms underlying this mode of migration.

To investigate the molecular drivers of CCM, we utilized MCF10A cells, a non-tumorigenic human mammary epithelial cell line that exhibits typical epithelial morphology, polarity, and genetic stability.^26^ These cells are widely used to model mammary epithelial biology and are highly responsive to microenvironmental cues, making them well suited for studying ligand-induced phenotypic transitions.^23,27–32^ Treatment with OSM induces CCM in MCF10A cells, establishing this system as a tractable model for dissecting the signaling pathways underlying CCM. ^23^ Our current study integrates live-cell imaging, transcriptomics, and functional assays to uncover the molecular programs driving CCM in MCF10A cells. We show that OSM activates a distinct transcriptional program in MCF10A cells, characterized by the upregulation of pathways associated with hypoxia and immune signaling. We identify hypoxia-inducible factor-1 (HIF1A) and signal transducer and activator of transcription 3 (STAT3) as central mediators of these responses and show that HIF1A contributes functionally to OSM-driven CCM.

## Results

### OSM induces collective migration in MCF10A epithelial cells

We previously reported that OSM induces changes in cell morphology and behavior, including collective cell migration (CCM).^23^ Motivated by this, we dissected the molecular mechanisms underlying OSM-mediated CCM by comparing responses of Oncostatin M (OSM), which signals via gp130/JAK-STAT,^24^ to epidermal growth factor (EGF), a canonical receptor tyrosine kinase (RTK) ligand^12^ and to interferon gamma (IFNG), another JAK-STAT activator.^33^ We chose IFNG as a comparator because it also activates JAK-STAT signaling but does not induce CCM. IFNG was delivered in combination with EGF, because unlike OSM, IFNG alone does not support proliferation of MCF10A cells.^23,34^

We treated MCF10A cells with OSM, EGF, or EGF + IFNG for 48 hours, and assessed their phenotypic responses with live-cell imaging and immunofluorescent microscopy. Staining with β-catenin at 48H revealed that EGF induces cells to form the characteristic cobblestone pattern typical of MCF10A epithelial sheets, while OSM treatment induces compact clusters of cells (**Figure 1A**).^29^ Live-cell imaging revealed that OSM-induced migration occurs as cohesive clusters, a hallmark of CCM (**Figure 1B**). This behavior was absent in both EGF and IFNG conditions.

**Figure 1:**
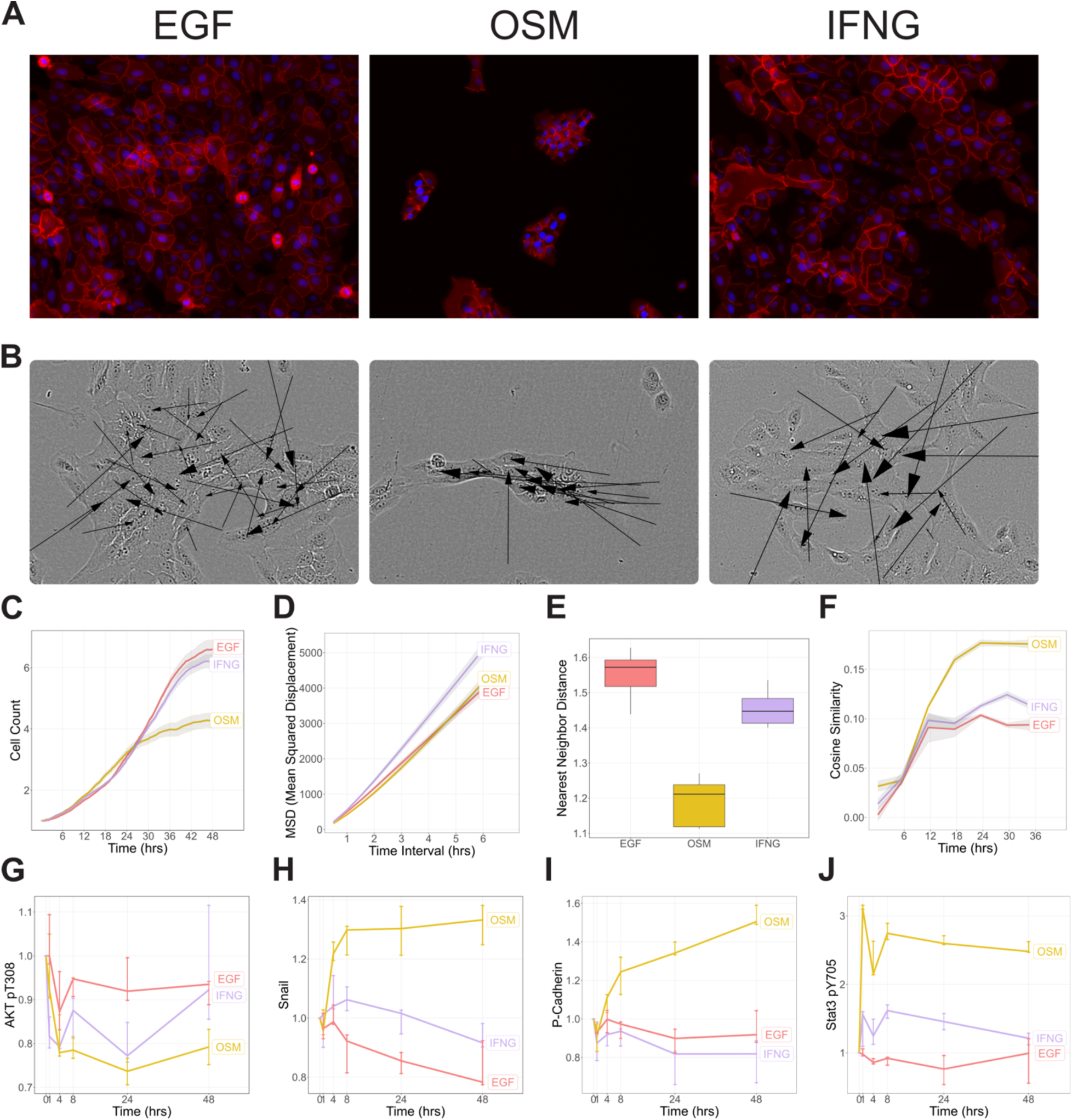
Treatment of MCF10A cells with OSM initiates cell clustering and collective cell migration. (A) MCF10A cells were treated with EGF, OSM, or IFNG for 48 hours, then stained for DAPI (blue) and β-Catenin (red). (B) Cell velocity vectors represent speed and direction of cell motility, derived from cell tracking data during 24-48 hours after ligand treatment. Arrow size is proportional to velocity. (C-F) Quantification of cell phenotype from live-cell imaging. Shaded region represents 95% confidence interval from 3 biological replicates (n=3). Boxplots in (E) depict interquartile range of nearest neighbor distance calculated for each cell. (G-J) Line plots show protein expression levels normalized to the T0 control for each treatment condition. Error bars represent the full range of measurements (n=3).

We quantified several metrics to capture key features of the behaviors observed using live-cell imaging data: cell count to estimate proliferation, mean squared displacement (MSD) to estimate motility, nearest neighbor distance to assess cell clustering, and cosine similarity of displacement vectors to assess collective migration (**Figure 1C-F**). Cell count was similar across conditions indicating minimal effects on proliferation (**Figure 1C, Supplemental Figure 1A**).

MSD revealed that IFNG treatment significantly increased motility of individual cells (p-value = .002), whereas OSM had no effect (**Figure 1D, Supplemental Figure 1A**). OSM significantly decreased nearest neighbor distance, indicating strong cell clustering (−22.3%, p-value = <.001), while IFNG showed only a modest reduction (−6.9%, p-value = .023) (**Figure 1E, Supplemental Figure 1A**). Cosine similarity, a measure of coordinated movement, was strongly elevated in OSM-treated cells, supporting coordinated, collective migration (p-value = <.001) (**Figure 1F**).

To identify molecular correlates of these phenotypic responses, we analyzed protein expression measured in publicly available reverse phase protein array (RPPA) data.^23^ EGF and EGF+IFNG induced stronger AKT activation than OSM, consistent with their more proliferative phenotypes (**Figure 1G**). RPPA further revealed that OSM uniquely upregulated Snail, an EMT-associated transcription factor (**Figure 1H**). This suggests that OSM-driven migration is accompanied by EMT-related transcriptional changes, while IFNG promotes motility without engaging EMT programs. OSM also uniquely upregulated P-cadherin and modulated other junctional proteins (Connexin-43, β-catenin, Merlin), consistent with increased cell-cell adhesion and clustering (**Figure 1I**). OSM and IFNG both activated STAT3 phosphorylation, though OSM induced a stronger response (**Figure 1J**). Despite this shared JAK-STAT activation, only OSM promoted CCM, whereas IFNG enhanced motility without inducing clustering or EMT marker upregulation. This suggests that although both ligands engage JAK-STAT signaling, OSM promotes a distinct molecular program associated with collective migration, whereas IFNG drives a more dispersed, single-cell migratory phenotype. This observation motivated us to further explore the contrast between OSM and IFNG as a tool for dissecting the molecular mechanisms underlying CCM in epithelial cells.

### Comparative network analysis identifies OSM-specific molecular regulators

To dissect the molecular mechanisms driving OSM-induced CCM, we analyzed publicly available profiling data from MCF10A cells treated with OSM or IFNG+EGF for 48 hours, including bulk RNA sequencing, RPPA, and cyclic immunofluorescence (cyclic IF).^23,35,36^ Our goal was to identify molecular signatures uniquely activated by OSM compared with IFNG, enabling us to distinguish regulators of collective migration from those mediating general JAK-STAT signaling.

We used CausalPath to infer network activity by integrating transcriptional and proteomic data, then visualized the network with Cytoscape^37,38^ (**Figure 2A, 2B Supplemental Figure 2**). We reasoned that molecular nodes showing the greatest differences in connectivity between OSM and IFNG conditions may act as key drivers of the OSM-induced CCM phenotype. To identify these candidates, we computed a rewiring score for each node, reflecting the number of edges that would need to be added or removed to shift its connectivity from one condition to the other.^39^ We weighted the rewiring metric by the log-fold change (LFC) induced by OSM for each node, prioritizing nodes that were both highly rewired and upregulated under OSM treatment.

**Figure 2:**
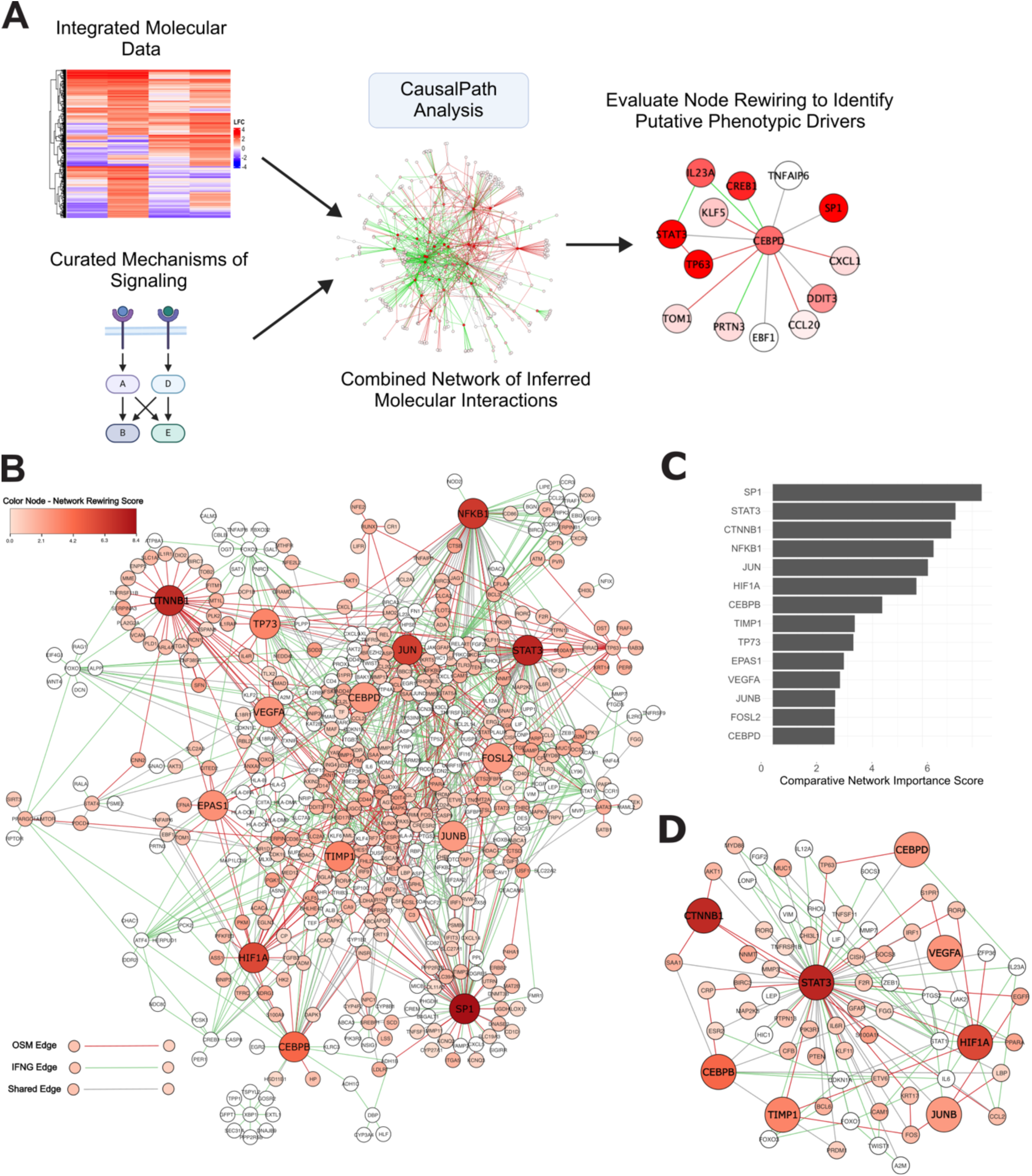
Comparative network analysis of omics data reveals putative drivers of OSM-induced CCM. (A) Workflow for the comparative network analysis to identify molecular subnetworks and nodes perturbed by OSM. Integrated molecular data collected from OSM and IFNG treated cells was analyzed using CausalPath. (B) Combined molecular network of OSM and IFNG activated nodes. The rewiring score between conditions is indicated by node color, and the top scoring nodes nominated for further investigation are indicated by increased size. (C) The top scoring nodes most rewired by OSM treatment, corresponding to the highlighted nodes in (B). (D) The majority (8/14) of top rewired nodes are centered around the STAT3 subnetwork.

STAT3 ranked second among nodes with the highest rewiring scores (**Figure 2C**). This demonstrates the value of a network-based approach: while STAT3 is activated by both treatments, its functional connectivity is distinct, suggesting unique functional roles. Eight of the fourteen top-rewired nodes were centered around the STAT3 subnetwork, consistent with the pivotal role of STAT3 in cytokine signaling and regulation of cellular phenotypes (**Figure 2D**).

Several transcription factors, including SP1, JUN family members, CTNNB1, TP73, and CEBP factors, were highly rewired, consistent with their roles in mediating cytokine signaling and regulation.^40–45^ Nodes associated with hypoxia signaling, such as HIF1A, VEGFA, TIMP1, and EPAS1, also showed high rewiring. HIF1A, a master regulator of hypoxia, is particularly notable as prior studies have indicated it can be upregulated by OSM in normal and cancerous cellular contexts.^46,47^ Hypoxia-related pathways contribute to modulating migratory phenotypes, including collective cell migration.^20,22,48^ These findings suggest a mechanistic link between OSM signaling and HIF1A-associated regulation of migratory behavior.

### Functional validation reveals causal regulators of OSM-induced CCM

We hypothesized that nodes uniquely rewired by OSM play a causal role in mediating CCM. To test this, we performed siRNA knockdown of the 14 nodes nominated in our network analysis In MCF10A cells under three experimental conditions: OSM, EGF, and IFNG+EGF treatments. We reasoned that knockdowns producing phenotypic effects specific to the OSM condition, rather than broadly affecting MCF10A cells across conditions, would identify regulators functionally critical for OSM-induced CCM. Cells were analyzed by live-cell imaging and immunofluorescence assessment at 48 hours post-treatment (**Figure 3A**), followed by quantitative assessment of cell count, MSD, nearest neighbor distance, and cosine similarity (**Figure 3B-E**).

**Figure 3:**
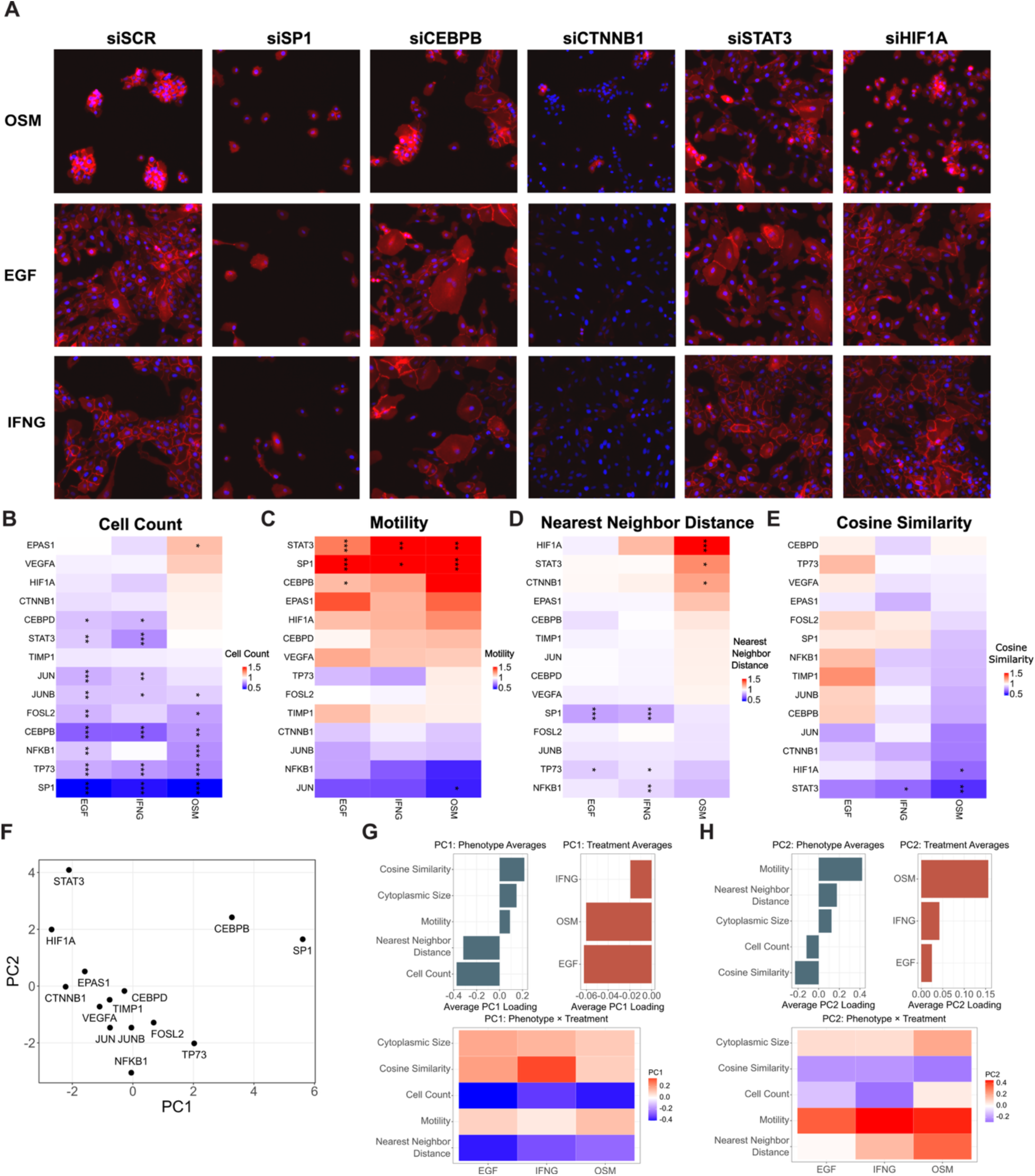
Quantification and visualization of siRNA experiment to identify drivers of OSM-induced CCM. (A) Representative immunofluorescent images from select knockdowns 48 hours after ligand treatment. (B-E) Quantification of cell phenotype induced by knockdown of nominated nodes (n=3). Phenotypic scores have been normalized to the siSCR condition for each ligand treatment. Statistical comparisons were performed using Student’s t-test. Significance is shown as follows: (* p-value < .05, ** p-value < .01, *** p-value < .001). (F) Principal component analysis of phenotypic metrics across all ligand treatments. (G-H) PC1 and PC2 loadings: Bar plots show average PC loadings for each phenotype and each treatment. The accompanying tile plot displays PC loadings across all phenotype x treatment combinations, with color indicating loading magnitude and direction.

To identify knockdowns that selectively perturbed OSM-induced behaviors, we performed principal component analysis (PCA) on the full set of phenotypic scores (**Figure 3F**). PC1 was negatively weighted by cell count and nearest neighbor distance, indicating that high PC1 scores reflect reduced proliferation and disrupted cell clustering (**Figure 3G**). Knockdowns of SP1 and CEBPB, which reduced cell number and altered cell shape across all ligand conditions, showed the strongest PC1 loadings. This PC appeared to capture general effects on cell viability or morphology, with no strong weighting from any specific ligand treatment, suggesting these regulators influence baseline epithelial functions rather than OSM-specific phenotypes (**Figure 3G**). In contrast, PC2 was positively weighted by motility and negatively weighted by cosine similarity, suggesting that high PC2 scores reflect increased individual cell motility and reduced collective migration behavior (**Figure 3H**). OSM treatment showed a strong positive weighting on PC2, indicating that knockdowns with high PC2 scores disproportionately disrupt OSM-induced phenotypes (**Figure 3H**). siRNAs targeting STAT3 and HIF1A were enriched along this axis, implicating them as central OSM-specific regulators (**Figure 3F**).

Consistent with their separation along PC2, knockdown of STAT3 and HIF1A increased nearest-neighbor distances and reduced cosine similarity under OSM treatment (**Figure 3D-E**). siSTAT3’s disruption of clustering reinforces its role as a master regulator of OSM-induced behaviors, consistent with its centrality in the OSM network. HIF1A knockdown specifically reduced cosine similarity and increased nearest neighbor distance under OSM treatment, suggesting a critical role for HIF1A signaling in OSM-induced CCM. HIF1A has been linked to migratory and invasive phenotypes through its regulation of cytoskeletal and extracellular matrix remodeling genes.^49,50^ The coordinated effects of HIF1A knockdown on clustering and cosine similarity support its function as a key regulator of OSM-induced CCM.

Additional OSM-specific knockdown phenotypes were observed. CTNNB1 knockdown increased nearest-neighbor distance only in the OSM condition, consistent with its role in maintaining cell-cell adhesion (**Figure 3D**).^51^ Knockdown of JUN specifically reduced motility in OSM-treated cells, suggesting it may promote the motile component of CCM (**Figure 3C**). EPAS1 (HIF2A) knockdown uniquely increased proliferation in the OSM context, an unexpected result given HIF2A’s reported role in promoting cell cycle progression, potentially revealing a context-dependent relationship between this transcription factor and proliferation (**Figure 3B**).^52^

Supplementary scatterplots comparing phenotypic scores across ligand conditions further supported these observations, particularly demonstrating the differential effects of HIF1A and STAT3 knockdowns on OSM-induced phenotypes (**Supplemental Figure 3A-B**). Comparing the magnitude of phenotypic changes under OSM versus IFNG revealed most knockdowns produced larger effects under OSM (ratios > 1). A binomial test confirmed this enrichment (p = 0.029), indicating that our comparative network approach successfully identified nodes that disproportionately impact phenotypes in the OSM context (**Supplemental Figure 3C**). We annotated the OSM-specific network with the phenotypic effects observed in our siRNA experiment, labeling each node based on its functional impact (**Supplemental Figure 4A-B**). This phenotype-annotated network provides a systems-level view of how distinct molecular regulators contribute to the diverse aspects of OSM-induced collective cell migration.

### HIF1A is a Key Mediator of OSM-Induced CCM

Our functional analyses identified HIF1A as a key driver of OSM-induced CCM, leading us to next dissect its mechanistic contributions. We returned to the RNA-seq and RPPA datasets used in our earlier analyses, which showed sustained upregulation of the HIF1A gene and protein over 48 hours of OSM treatment, an effect absent under EGF or IFNG (**Figure 4A, Supplemental Figure 5A**). To identify the transcriptional programs HIF1A activates in OSM-treated cells, we generated MCF10A cells with stable shRNA-mediated HIF1A knockdown (**Supplemental Figure 5B**). We treated shHIF1A and shSCR control cells with OSM, EGF, or IFNG and performed single-cell RNA-seq at 24 hours. Assessment of HIF1A gene expression confirmed the knocked down in shHIF1A cells (**Supplemental Figure 5C**).

**Figure 4:**
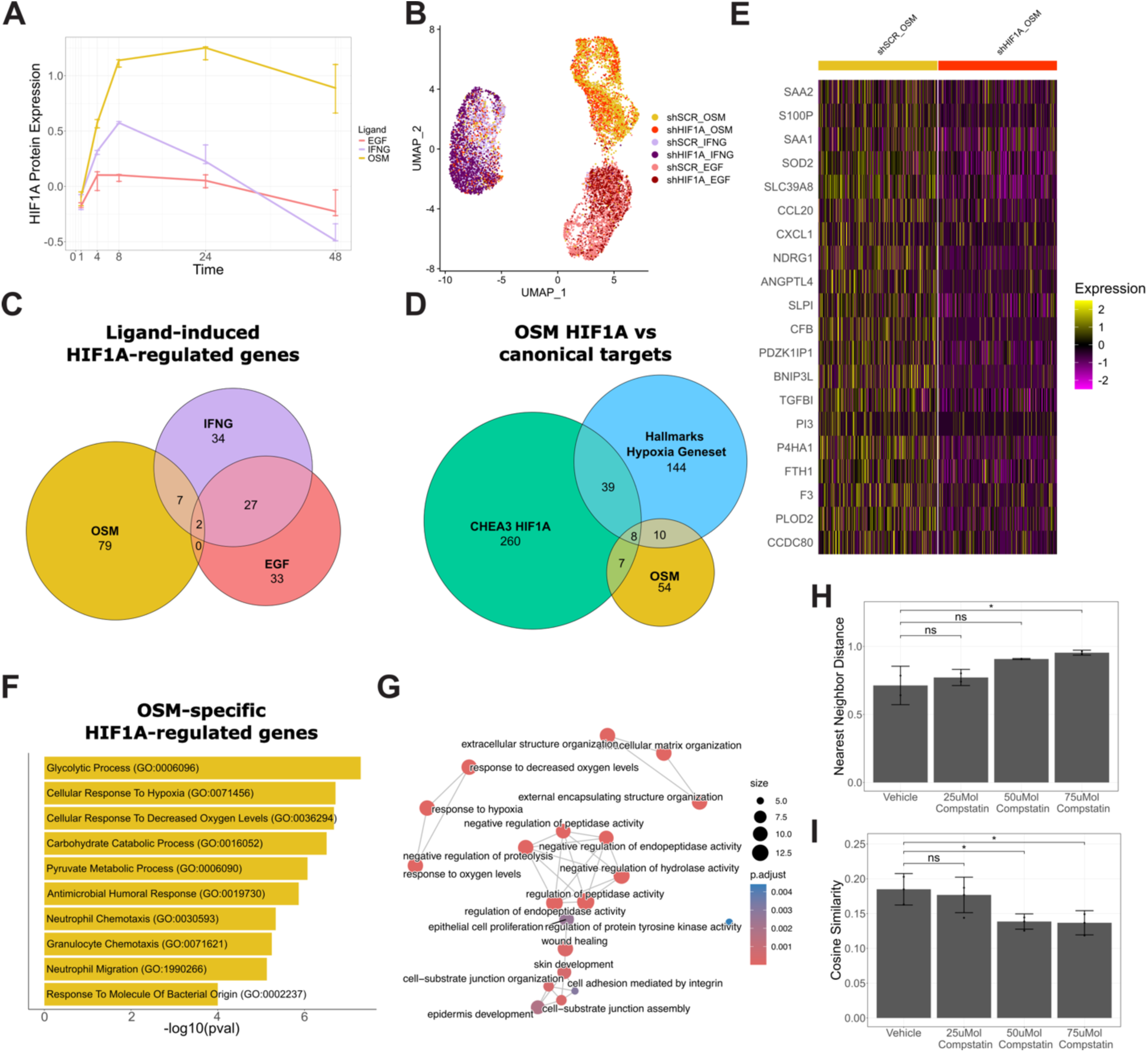
scRNAseq reveals HIF1A transcriptional program driving OSM-induced CCM. (A) OSM uniquely upregulates the HIF1A protein over 48 hours of ligand treatment. Error bars represent standard deviation (n=3). (B) UMAP projection of scRNAseq data indicates that transcriptional signals separate primarily by ligand treatment. (C) We identified HIF1A-regulated genes for each ligand condition by performing differential gene expression analysis between shSCR vs shHIF1A. Venn diagram represents a comparison of HIF1A-regulated gene sets under each treatment. (D) Comparison of unique OSM-HIF1A regulated genes to literature derived gene sets representing hypoxia and HIF1A transcription factor activity. (E) Relative expression of the top 20 differentially expressed unique OSM-HIF1A regulated genes. (F) We performed gene set enrichment analysis to identify Gene Ontology terms enriched in OSM-HIF1A regulated genes. Top ten most enriched gene sets are shown. Multiple comparison adjustments were made using the Benjamini-Hochberg method. (G) Enrichment map of Gene Ontology terms enriched in shSCR cells under OSM treatment. (H) Inhibition of complement signaling increases nearest neighbor distance in OSM treated cells and reduces cell clustering. Error bars represent the 95% confidence interval (n=3). Statistical comparisons were conducted using Dunnett’s test, and significance was defined as p-value < .05. (I) Compstatin treatment reduces cosine similarity in OSM treated cells and inhibits CCM. Error bars represent the 95% confidence interval (n=3). Statistical comparisons were conducted using Dunnett’s test and significance was defined as p-value < .05.

UMAP projection of scRNA-seq data showed that cells clustered primarily by ligand treatment, regardless of HIF1A knockdown status (**Figure 4B**). To isolate transcriptional changes driven specifically by OSM-activated HIF1A, we compared differentially expressed genes in OSM-treated shHIF1A versus shSCR cells and excluded genes regulated by HIF1A under EGF or IFNG treatment. This analysis revealed 79 genes uniquely regulated by HIF1A in the OSM condition. (**Figure 4C**).

We more deeply examined the 79 OSM-specific HIF1A-regulated genes to prioritize candidate drivers of CCM. First, we asked whether these genes have already been linked to HIF1A and hypoxia signaling by comparison to Hallmark and ChEA3 gene sets; this confirmed overlap with 25 canonical hypoxia targets including NDRG1 and CA12 (**Figure 4D, Supplemental Figure 5D-E**).^53–56^ However, OSM-HIF1A also uniquely upregulated 54 additional genes not typically associated with hypoxia or canonical HIF1A transcriptional regulation, suggesting the activation of a specialized transcriptional program in the OSM context. These genes exhibited consistent upregulation in OSM-treated cells compared to other conditions (**Figure 4E**). Second, gene set enrichment analysis (GSEA) showed that these 79 genes were enriched for glycolysis and metabolic reprogramming pathways, including “glycolytic process”, “carbohydrate catabolic process”, and “pyruvate metabolic process” (**Figure 4F**).^57^ We also observed enrichment of hypoxia-related gene sets, including “cellular response to hypoxia” and “cellular response to decreased oxygen levels.” These findings align with HIF1A’s well-documented role in promoting glycolysis and mediating the cellular response to hypoxia, confirming that HIF1A retains aspects of its canonical functionality in the OSM condition.^58^

In addition to metabolic and hypoxia-related processes, we observed enrichment of immune-related pathways, particularly those involving neutrophil migration and chemotaxis. Because neutrophils engage in collective migration, we hypothesized that this transcriptional signature may relate to the observed OSM-induced CCM phenotype.^59–61^ Closer examination of the enrichment map linking these gene sets revealed HIF1A-regulated genes involved in complement signaling and regulation of peptidase activity (**Figure 4G, Supplemental Figure 5E**), a pathway known to support cell adhesion and collective migration in other biological contexts.^62–64^

To test whether complement signaling functionally contributes to OSM-induced CCM, we treated cells with OSM and Compstatin, an inhibitor of C3, the central hub of complement activation that drives downstream effector pathways.^65^ Live-cell imaging showed that Compstatin treatment reduced clustering (**Figure 4H** (25mmol p-value = .630, 50mmol p-value = .056, 75mmol p-value = .028) and cosine similarity (25mmol p-value = .646, 50mmol p-value = .027, 75mmol p-value = .023) in a dose-dependent manner, mirroring phenotypes observed with HIF1A knockdown (**Figure 4I, Supplemental Movie 1,2**).These findings establish a mechanistic link between OSM signaling, HIF1A activation, complement signaling, and CCM. The HIF1A-dependent transcriptional program induced by OSM encompasses canonical hypoxia targets as well as genes involved in immune and complement-related pathways, consistent with a broader regulatory role for HIF1A in epithelial collective migration.

### OSMR and its downstream transcriptional programs are active in human breast tumors and associated with poor prognosis

Given the role of CCM in cancer invasion and metastasis, we hypothesized that OSM signaling and its downstream effectors contribute to poor outcomes in breast cancer patients. High OSMR expression was significantly associated with reduced overall survival in patients with PAM50 classified basal breast cancers (log-rank P = 0.034) (**Figure 5A**).^71,70^ To identify the cell types driving or responding to OSM signaling, we analyzed publicly available single-cell RNA-sequencing data from clinical breast cancer samples.^66^ OSM expression was primarily restricted to tumor-infiltrating myeloid cells (**Figure 5C**), whereas OSMR expression was enriched in multiple non-myeloid cell types, including a subset of cancer epithelial cells, endothelial cells, fibroblasts, and perivascular cells (**Figure 5D**). These findings suggest that OSM signaling is active in the tumor microenvironment and may influence both malignant and stromal components, potentially driving aggressive tumor behavior.

**Figure 5:**
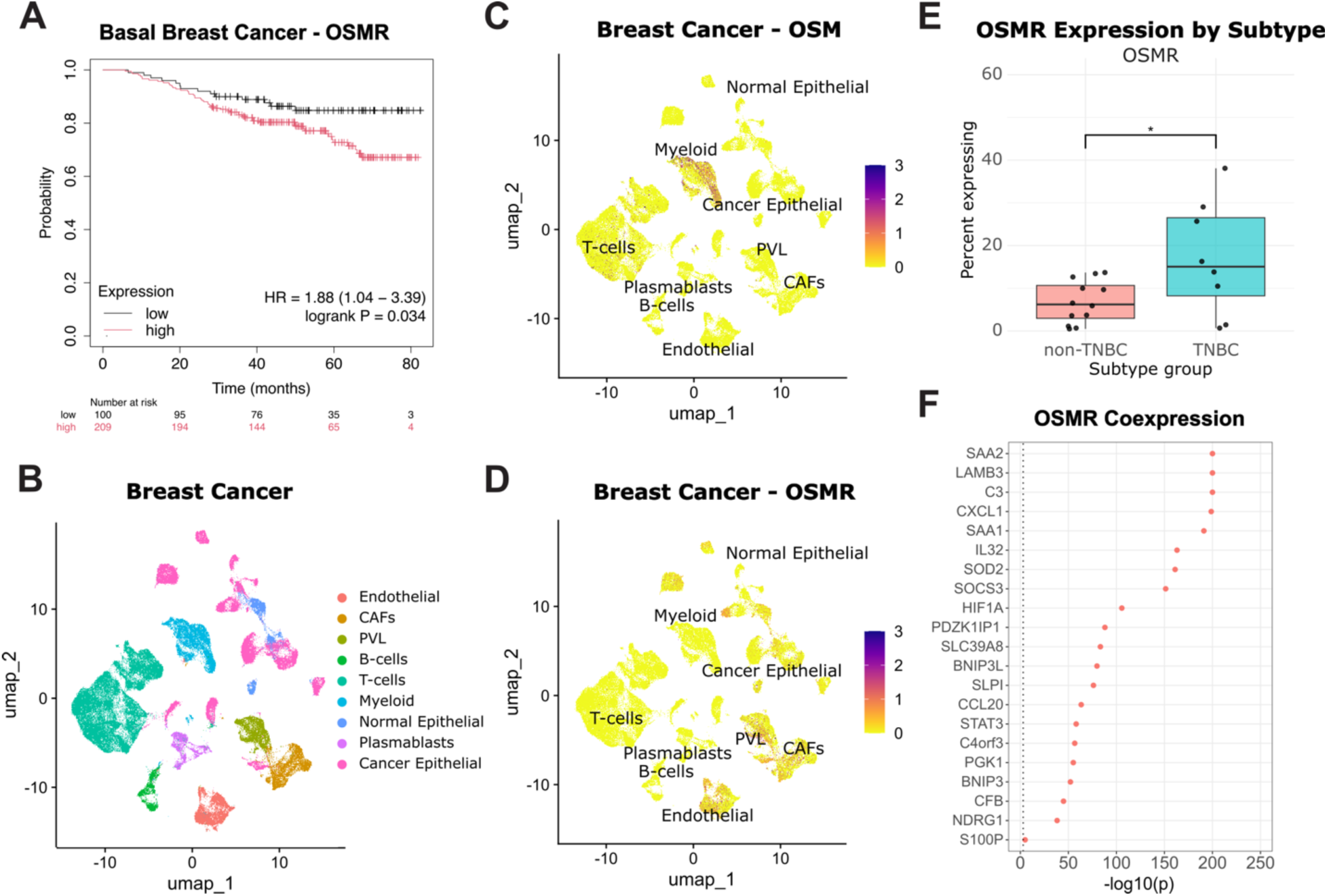
The OSM–HIF1A signaling axis is active in basal-like breast cancer and associated with poor prognosis. (A) Kaplan–Meier survival analysis of patients with basal-like breast cancer stratified by OSMR expression (TCGA-BRCA, n = 309). High OSMR expression is associated with reduced overall survival (log-rank P = 0.034). (B) UMAP of single-cell RNA-seq data from human breast tumors annotated by cell type. (C) UMAP showing OSM is primarily expressed by myeloid cells in breast cancer tumors. (D) UMAP showing OSMR expression in stromal and epithelial tumor compartments, including epithelial cancer cells. (E) Boxplot showing the percentage of epithelial cells expressing OSMR in individual patients, comparing triple-negative breast cancer (TNBC) to non-TNBC tumors. TNBC tumors have a higher proportion of OSMR-positive epithelial cells (* p-value < .05). (F) COTAN analysis of OSMR coexpression within cancer epithelial cells shows robust coexpression of mechanistic signature. Dotted line indicates significance threshold (p-value < .01).

To further explore the role of OSMR signaling in cancer epithelial cells, we stratified tumors by molecular subtype. Epithelial cells from triple-negative breast cancer (TNBC) tumors were more likely to express OSMR (p-value = .0287) and C3 (p-value = .00149) compared to other subtypes (**Figure 5E, Supplemental Figure 6A**). Within each subtype, the percentage of epithelial cells expressing OSMR varied significantly by patient, reflecting inter-patient heterogeneity. (**Supplemental Figure 6B**). Finally, we assessed whether the components of the OSM–HIF1A transcriptional program are coexpressed within individual cancer epithelial cells using COTAN analysis.^67^ This revealed that all genes in the signature, including HIF1A and C3, are significantly coexpressed in malignant epithelial cells (**Figure 5G**).

Together, these results demonstrate that the OSM–HIF1A signaling axis and its downstream transcriptional programs are active in basal-like breast tumors. The enrichment of OSMR expression and expression of its target genes in epithelial cancer cells, along with their association with poor survival, supports the potential clinical relevance of this cytokine-driven collective migration mechanism. These observations extend our findings to human disease and suggest that targeting the OSM-HIF1A axis could have therapeutic value for the subset of TNBC patients with high OSMR expression.

## Discussion

Cell migration is a key process in various physiological and pathological contexts, including tissue development, wound healing, and cancer metastasis. CCM, in which groups of cells move together as a cohesive unit, plays a critical role in maintaining tissue architecture and facilitating invasion during cancer metastasis.^1^ The mechanisms driving CCM are complex, involving interactions between signaling pathways, cytoskeletal dynamics, and cell-cell communication. In this study, we explored how the cytokine OSM drives CCM in mammary epithelial cells. Through network analysis and experimental validation, we identified HIF1A and STAT3 as central regulators of OSM-induced CCM.

Our results demonstrate that OSM induces a unique transcriptional program that drives CCM, with HIF1A and STAT3 acting as key regulators. Using siRNA, we mapped the functional role of these nodes in modulating phenotypic responses. HIF1A knockdown significantly impaired OSM-induced CCM, confirming its central role in this process. Similarly, STAT3, a canonical mediator of cytokine signaling, was required for the induction of OSM-dependent clustering and migration.^68^ Together, these findings suggest that OSM promotes a coordinated cellular response that integrates hypoxic and inflammatory cues to drive collective migration.

HIF1A is classically known for its role in regulating cellular responses to hypoxia, including metabolic reprogramming, survival, and motility.^69^ More recently, HIF1A has been implicated in promoting partial epithelial-to-mesenchymal transition (EMT), a transcriptional and phenotypic program that enhances cellular plasticity, motility, and invasiveness while preserving epithelial characteristics such as cell-cell junctions^20,70^. Our data support this model; OSM treatment upregulated HIF1A expression and activated downstream gene programs that resulted in cells retaining intercellular junctions while acquiring migratory capacity, which are hallmarks of partial EMT.^71,72^ These results reinforce the connection between inflammatory signaling and partial EMT ^73–75^. These findings may be particularly important in cancer biology, where partial EMT has been increasingly recognized as a driver of collective invasion and metastasis.^76–78^ Unlike full EMT, which involves complete loss of epithelial identity, partial EMT states may allow cells to migrate collectively while maintaining cooperative interactions and intercellular signaling that promotes cell survival.^79^ Our data extend this framework by implicating OSM, through HIF1A, as an inducer of this hybrid migratory state.

An additional finding in our study was the enrichment of complement-related immune pathways in HIF1A-regulated genes. This prompted us to investigate whether complement activation functionally contributes to OSM-induced migration. Using Compstatin, a C3-specific inhibitor, we observed a marked reduction in both clustering and motility, mirroring the effects of HIF1A knockdown. This finding supports a model in which complement signaling acts downstream of HIF1A to mediate cellular behaviors necessary for CCM. The identification of complement as an effector pathway adds to our understanding of how immune and hypoxic signals converge to drive collective migration and suggests that targeting complement signaling may offer an approach to modulate these behaviors in disease contexts. However, the precise mechanism by which complement activity regulates CCM remains unclear. Possible modes of action include modulation of coattractive autocrine/paracrine signaling through complement receptors,^63^ activation of chemotaxis signals and their downstream effectors,^62^ or indirect effects on integrin regulation and cell–cell adhesion.^64^ Elucidating these possibilities will require future work to disentangle the direct versus secondary consequences of complement inhibition. In addition, gene set enrichment analysis revealed a strong upregulation of glycolysis-related genes, further linking HIF1A to metabolic reprogramming in OSM-treated cells. While glycolytic shifts are a well-known feature of the hypoxic response, our data raise the possibility that OSM-induced HIF1A activity may contribute to broader metabolic changes in tumors.^80,81^ These shifts could reflect a reallocation of resources to meet the energetic and biosynthetic demands of collective cell migration.^82,83^ Moreover, the induction of a glycolytic program suggests that OSM signaling, via HIF1A, may act as a broader regulator of cellular state transitions, promoting not only motility but also metabolic adaptations.

Motivated by our in vitro findings, we investigated the relevance of this pathway in breast cancer by analyzing publicly available scRNA-seq data.^66^ We found that OSM is primarily expressed by tumor-infiltrating myeloid cells, whereas OSMR, the cognate receptor, is expressed by a subset of epithelial tumor cells. Notably, OSMR expression was highest in triple-negative breast cancer (TNBC) tumors, a subtype molecularly aligned with basal breast cancer and associated with poor prognosis.^84^ This suggests that OSM-driven signaling may be active in aggressive cancer subtypes. Supporting this, we found that OSMR expression is negatively associated with survival in patients with basal breast cancer. Our analysis also revealed significant coexpression among OSMR, HIF1A, and our HIF1A-induced gene signature in malignant breast cancer epithelial cells. These findings reinforce the idea that the OSM-HIF1A-complement axis constitutes a functionally active pathway in vivo. While we have not directly assessed the role of the OSM axis in primary versus metastatic disease, the link between CCM and metastatic dissemination^79^ raises the possibility that OSM-induced CCM could be an important driver of metastasis. In this context, the association between high OSMR expression and poor survival further supports the potential clinical significance of this pathway in disease progression.

While our study provides a comprehensive characterization of OSM-induced CCM in MCF10A cells, it has several limitations. Most notably, our findings are based on a non-transformed mammary epithelial cell line. Although this model offers a controlled environment to study cytokine responses, future efforts could extend these observations to cancer-derived organoids,^85^ xenografts,^86^ or patient samples^87^ to evaluate how OSM-driven CCM operates in diverse tumor microenvironments. In addition, while we focused on transcriptional changes, CCM is also regulated by post-translational modifications,^88,89^ cytoskeletal rearrangements,^90^ and dynamic cell-cell interactions,^91^ which could be further explored with advanced imaging approaches. Additionally, our analysis focused primarily on the transcriptional consequences of HIF1A activation, yet HIF1A also modulates various proteomic signaling pathways that could influence CCM.^1^ Pathways like mTOR signaling, integrin-mediated adhesion, and autophagy, which are also regulated by HIF1A, may contribute to CCM by altering cellular motility, adhesion, and metabolic support for migration^49,92,93^. Further investigation into how HIF1A coordinates these signaling pathways will provide a more comprehensive understanding of its role in CCM.

In conclusion, our findings implicate HIF1A as a key mediator linking inflammatory cytokine signaling and CCM, processes relevant to cancer progression. We also observed induction of SNAI, suggesting that OSM engages elements of partial EMT, another mechanism associated with metastatic potential.^20,78^ Consistent with this, we observed that OSMR expression negatively correlates with survival in basal breast cancer, and that breast epithelial cells frequently coexpress OSMR, HIF1A, and our hypoxia-associated gene signature. Our results are consistent with prior work implicating OSM in cancer progression and provide a foundation for future efforts to target these pathways in cancer metastasis and tissue remodeling.

## Methods

### MCF10A Cell Culture

Cell culture and ligand perturbation experiments were carried out according to the methods described by Gross et al. 2022.^23^ Briefly, for routine cell growth and passaging, MCF10A cells were maintained in a growth medium consisting of DMEM/F12 (Invitrogen #11330-032), 5% horse serum (Sigma #H1138), 20 ng/ml EGF (R&D Systems #236-EG), 0.5 µg/ml hydrocortisone (Sigma #H-4001), 100 ng/ml cholera toxin (Sigma #C8052), 10 µg/ml insulin (Sigma #I9278), and 1% Pen/Strep (Invitrogen #15070-063). For perturbation studies, a growth factor-free medium was used, which included DMEM/F12, 5% horse serum, 0.5 µg/ml hydrocortisone, 100 ng/ml cholera toxin, and 1% Pen/Strep.

MCF10A cells were cultured until they reached 50-80% confluence in growth medium, followed by detachment using 0.05% trypsin-EDTA (Thermo Fisher Scientific #25300-054). After detachment, 6,000 cells were seeded into collagen-1 (Cultrex #3442-050-01) coated 24-well plates (Thermo Fisher Scientific #267062) with growth medium. Six hours later, the cells were washed with PBS, and growth factor-free medium was added. The cells were then incubated for 18 hours in the new medium. Afterward, cells were treated with either 10 ng/ml EGF (R&D Systems #236-EG), 10 ng/ml OSM (R&D Systems #8475-OM), or 20 ng/ml IFNG (R&D Systems #258-IF) + 10 ng/mL EGF.

### Live-Cell Imaging

Live-cell imaging was conducted using the Incucyte S3 microscope (Essen BioScience, #4647), with images captured every 30 minutes for a duration of up to 48 hours. All experiments were performed using three biological replicates. The resulting live-cell image stacks were processed by first registering them with a custom Fiji script and then segmented using CellPose v3.01.^94,95^ Image tracking was performed using the Baxter Algorithms pipeline.^96^

Subsequent analysis of the cell tracking data was conducted within RStudio.^97^ Cell counts were calculated by determining the number of cells per field, normalized by the initial T0 count for each field. Nearest neighbor distances were computed by measuring the Euclidean distances in pixels from the centroid of each cell to the centroid of the second nearest cell within the imaging field. To adjust for variations in cell count, the average nearest neighbor distances for each image were normalized by the expected mean distance to the nearest neighboring cell under a random cell distribution model.^98^ Cell motility was assessed by excluding tracks with jumps greater than 200 pixels in a 30-minute interval, as these tracks likely represent tracking errors. Motility was quantified as the slope of the mean squared displacement (MSD) over time intervals ranging from 30 minutes to 6 hours.^99^ This slope was determined by constructing a linear model comparing MSD to time intervals, and is proportional to the diffusion coefficient associated with Brownian motion.^99^ Finally, collective cell migration was estimated by calculating the cosine similarity between displacement vectors of the 10 nearest cells every 30 minutes. Statistical comparisons of phenotypic scores between ligand conditions was performed using Dunnett’s test, with EGF serving as the control.^100^

### Immunoflourescent Imaging

For all immunofluorescent experiments, cells were fixed in 4% formaldehyde after 48 hours of ligand treatment. Cells were blocked for 1 hour in a PBS solution (5% Normal Goat Serum (Cell Signaling #5425), 0.3% Triton X (Thermo Fisher Scientific #85111), then incubated overnight with the primary conjugated antibody (Cell Signaling #2677) at a 1:500 dilution. Cells were washed thoroughly then counterstained with 0.5 mg / mL DAPI (PromoKine PD-CA707-40043). Imaging was performed using the InCell 6000 (GE Healthcare).

### RNAseq, RPPA Data, and Cyclic IF

The bulk RNA sequencing (RNAseq), reverse phase protein array (RPPA) data, and cyclic immunofluorescence (cycIF) data after ligand treatments were generated in a prior investigation.^23^ For detailed information on the methodologies used for data generation, normalization, and integration, please refer to the original publication.

### Causal Pathway Analysis and Network Analysis

Causal Pathway was utilized to construct gene and protein interaction networks.^37^ The input to the algorithm was the integrated features from RNAseq, RPPA, and cyclic IF datasets. Data from proteomic assays were favored for features shared between assays. Only features with a log-fold change (LFC) of 1 or greater compared to the time zero control were included in the analysis. The generated networks were subsequently integrated and analyzed in Cytoscape.^38^

To identify gene/protein nodes for experimental testing, a comparative analysis was performed between the IFNG and OSM networks. Each node was assigned a score based on network rewiring, when comparing the OSM network to the IFNG network using Dynet.^39^ This score was then scaled according to the LFC in the OSM condition. The fourteen nodes with the highest comparative network importance scores were selected for experimental validation.

### siRNA Screen and Drug Experiments

Reverse transfection of MCF10A cells was performed according to DharmaFECT Transfection vendor protocols. Cells were seeded in collagen coated 24-well plates with growth media as previously described (see methods: MCF10A cell culture). After 6 hours in culture, media was replaced with 25 nM siRNA and 0.5 mL Lipofectamine™ RNAiMAX Transfection Reagent (Thermo Fisher Scientific #13778075) per well in antibiotic free media. After 48 hours of transfection, cells were washed thoroughly with PBS and replaced with assay media and indicated ligand treatments. The siRNA screen was run in triplicate. A full list of siRNAs used in the study is included (**Supplemental Table 1**). Immediately after ligand treatment, live-cell imaging and phenotypic quantification was performed as described previously (see Methods: Live-Cell Imaging). Statistical comparisons between phenotypic scores were performed by pairwise Student’s t-tests between knockdown conditions and the siSCR condition for each ligand treatment.^101^ Principal Component Analysis (PCA) was performed on phenotype quantifications using the prcomp function from the stat R package (3.6.2).^97^

For drug experiments, cells were cultured and plated as previously described (see Methods: MCF10A cell culture). Concurrent to ligand treatment, cells were incubated with either vehicle, or 25 mMol - 75 mMol Compstatin (Selleckchem #S8522). Statistical comparisons between treated and untreated conditions were conducted using Dunnett’s test.^100^ Significance was defined as p-value < .05.

### shRNA Integration

Plasmid constructs for non-targeting SMARTvector Lentiviral shRNA and shRNA sequences targeting HIF1A were purchased from Horizon Discovery (Horizon Discovery # VSC11287, #3SH11243-02EG3091). Expression of the shRNA was driven by the EF1A promoter and contained Puromycin resistance and TurboGFP elements to ensure proper integration. *E. Coli* containing the plasmid constructs were inoculated into LB broth medium containing 100 μg/mL carbenicillin and incubated at 37C for 18 hours with shaking. Plasmids were extracted using the HiSpeed Plasmid Midi Kit (Qiagen #12643).

Plasmids were then packaged into lentiviral particles by transfecting 293T human embryonic kidney cells (ATCC #CRL-3216) with the shRNA plasmid construct and lentiviral package plasmids (Addgene #8454, #12263) using LF2000 transfection reagent (Invitrogen #11668019). Virus containing media was collected through a 0.45 mm low-protein binding filter (Sterlitech #PES4547100). MCF10A cells were then transduced with virus containing media and 10 mg/mL polybrene. After a four-hour transduction, media was replaced with growth media, and shRNA containing cells were selected with 0.5 mg/mL puromycin. Expression of shRNA was confirmed using fluorescent imaging of GFP on the Incucyte S3 microscope (Essen BioScience #4647).

### Immunoblotting

HIF1A was detected in cells transduced with shRNA through immunoblotting. After 24 hours of treatment with OSM EGF, or CoCl2 cells were lysed in hot Laemmli buffer (Santa Cruz Biotechnology #sc-286962). Cell lysates were boiled at 95C for 10 minutes, then total protein concentration was determined and normalized through BCA assay (Thermo Fisher Scientific #23225). 25 mL of cell lysate and HIF1A lysate control (Novus Biologicals #NBP2-04440) was loaded onto gels (Invitrogen #NP0321BOX) and proteins were separated at 200V for 45 minutes then transferred at 30V overnight at 4C. After transfer the membrane was blotted with TBST and 5% dry milk for 1 hour, then probed for HIF1A at a primary antibody concentration of 1:1000 (BD Biosciences #610959) and secondary antibody 1:10000 (Jackson Immunoresearch #715-035-150). Chemiluminescence was imaged on the Alpha Innotech FluorChem Imaging System (#22424).

### scRNA-seq Library Preparation and Sequencing

All cell lines and ligand conditions were multiplexed using Hashtag Oligonucleotide barcoding (TotalSeq-A, Biolegend #399907) according to the vendor’s recommendations. The mRNA library was created using the Chromium Single Cell 3ʹ GEM, Library & Gel Bead Kit v3 (10X Genomics #1000092) and then sequenced using an Illumina NovaSeq with a depth of 800 million reads. Short read sequencing assays were performed by the OHSU Massively Parallel Sequencing Shared Resource.

### scRNA-seq Data Processing and Analysis

Raw files were converted to FASTQ format with bcl2fastq (version 2.20.0.445). Cellranger count was used to align reads to the GRCh38 transcriptome (GCF_000001405.40). The R package deMULTIplex was used to demultiplex the hash-tagged samples and assign cell lines and treatments to cells.^102^ Seurat (4.0.5) was used to perform variable feature identification, dimensionality reduction, unsupervised clustering, visualization, and differential gene expression.^103^ Differential expression analysis was performed using the FindMarkers function of Seurat with default parameters. GSEA was performed with the R package clusterProfiler on Gene Ontology Biological Process gene sets using significantly upregulated genes compared to the shSCR control (LFC > 0.5, p-value < .05).^104^

### Survival Analysis

To assess the prognostic significance of OSMR and OSM-HIF1A target gene expression, we utilized the Kaplan–Meier Plotter (KmPlot) web tool, which integrates public RNA sequencing data from breast cancer patient cohorts.^105,106^ Patients were stratified into two groups based on gene expression levels using the tool’s automatic cutoff selection algorithm, and overall survival was compared using the log-rank test.

### Breast Cancer Single-Cell RNA Sequencing Analysis

To investigate OSM pathway activity within the tumor microenvironment, we analyzed a publicly available single-cell RNA sequencing atlas of 26 primary human breast tumors.^66^ Processed expression matrices were analyzed using Seurat (4.0.5).^103^ The pipeline included variable feature identification, dimensionality reduction, unsupervised clustering, visualization, and differential gene expression analysis. Differential expression was performed using Seurat’s FindMarkers function, applying a p-value threshold of < 0.01 and a log2 fold-change cutoff > 1. Cell type and breast cancer subtype annotations were obtained from the original publication. To quantify transcriptional coordination of OSM-HIF1A genes in malignant epithelial cells, we calculated coexpression scores and estimated p-values using the COTAN package.^67^ Genes with a coexpression p-value < 0.001 were considered significantly coexpressed.

## Supporting information

MovieS1 - Compstatin50mM

MovieS2 - Vehicle

## Data Availability

### Lead contact

Further information and requests should be directed to the lead contact, Laura M Heiser.

Raw live-cell imaging and IF imaging data of MCF10A cells treated with siRNA knockdown and OSM, EGF, or IFNG are available Zenodo: 10.5281/zenodo.17101147, 10.5281/zenodo.17122918, 10.5281/zenodo.17136555, 10.5281/zenodo.17136574. Single-cell RNA-seq data generated in this study have been deposited in the Gene Expression Omnibus (GEO) under accession number: GSE307221.

### Code Availability

All analysis scripts, along with a complete list of R packages and versions used, are available at: https://github.com/mcleania/MCF10A_OSM_HIF1A. Any additional information required to reanalyze the data reported in this paper is available from the lead contact upon request.

## Acknowledgments

This work was supported by NIH research grants U54-CA209988, U54 HG008100, the Anna Fuller Fund, and the Jayne Koskinas Ted Giovanis Foundation for Health and Policy (L.M.H.). OHSU Massively Parallel Sequencing Shared Resource receives support from the OHSU Knight Cancer Institute NCI Cancer Center Support Grant P30CA069533. The authors acknowledge Lauren Kronebusch for help with manuscript editing. Cartoons created with Biorender.com. The Jayne Koskinas Ted Giovanis Foundation for Health and Policy is a private foundation committed to critical funding of cancer research. The opinions, findings, conclusions or recommendations expressed in this material are those of the author(s) and not necessarily those of the Jayne Koskinas Ted Giovanis Foundation for Health and Policy or its respective directors, officers, or staff. The research reported in this publication used computational infrastructure supported by the Office of Research Infrastructure Programs, Office of the Director, of the National Institutes of Health under Award Number S10OD034224. The content is solely the responsibility of the authors and does not necessarily represent the official views of the National Institutes of Health.

## Author Contributions

Conceptualization: LMH, ICM, and SMG; Investigation: ICM, SMG, TL; Methodology: LMH, ICM, MD; Formal Analysis: ICM, DSD, MAD; Visualization: ICM; Supervision: LMH; Writing—Original Draft: ICM, Writing—Review and Editing: LMH, ICM, and DSD.

## Supplementary Information

Supplemental Table 1 – siRNA catalogue

Supplemental Video 1,2 – Live cell imaging of vehicle or complement inhibition on OSM treated MCF10A cells

