## Supplemental Figures for "Oncostatin M orchestrates collective epithelial migration via HIF1A activation"

A

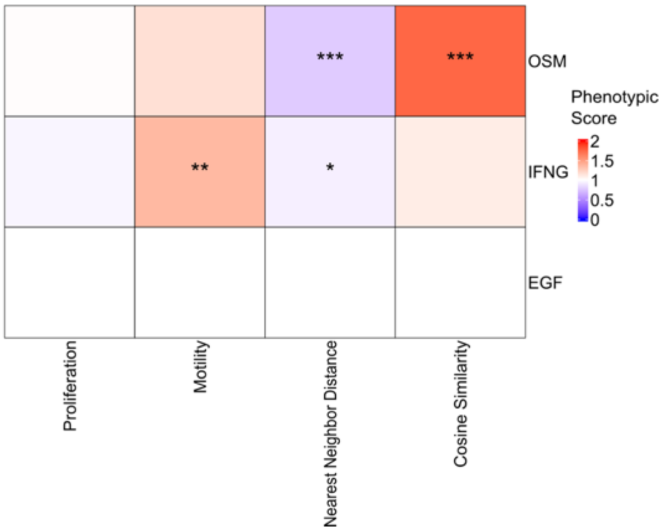


**Supplemental Figure 1: Statistical comparison of phenotypic metrics.**

(A) Statistical comparison of the quantifications of cellular proliferation, motility, nearest neighbor distance, and cosine similarity. Phenotypic scores have been normalized to the EGF condition. Dunnett's test was employed to compare phenotypic scores to the EGF control (n=3). Significance is shown as follows: (* p-value < .05, ** p-value < .01, *** p-value < .001).

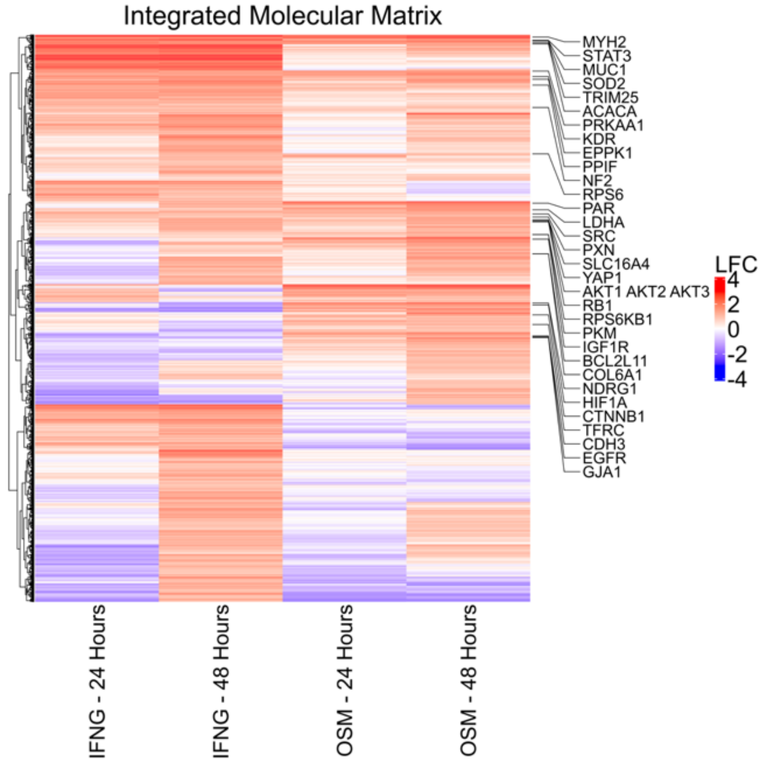


**Supplemental Figure 2: Integrated and filtered molecular data collected from OSM and IFNG treated cells**

Proteomic (RPPA, CycIF) and transcriptional (RNAseq) data was collected from MCF10A cells treated with OSM or IFNG for 24 and 48 hours (n=3). The data from each assay was normalized to the Time 0 control, filtered for differentially expressed features (LFC > 1, p-value < .05 adjusted by FDR) and then integrated into a combined dataset. Proteomic features from the RPPA assay are labelled.

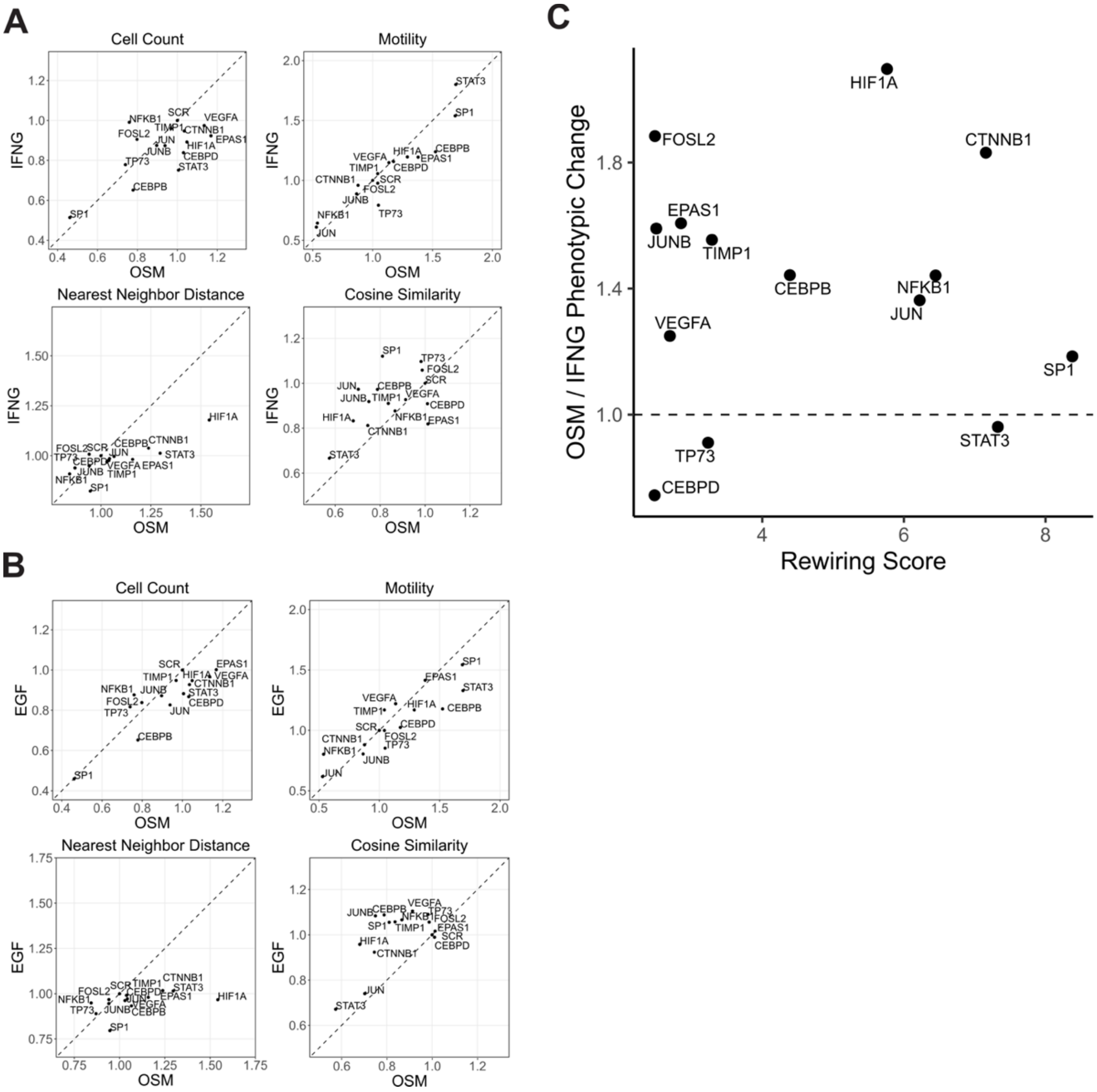

**Supplemental Figure 3: Comparison of phenotypic effects of knockdown between ligand conditions**

(A-B) Comparison of the phenotypic effects of knockdown on OSM and (A) IFNG or (B) EGF treated cells. (C) Scatterplot comparing the rewiring score to the overall magnitude of phenotypic change for each knockdown relative to the scramble control. Nominated nodes were more likely to exert greater effects under OSM treatment compared to IFNG (binomial test, p = 0.029).

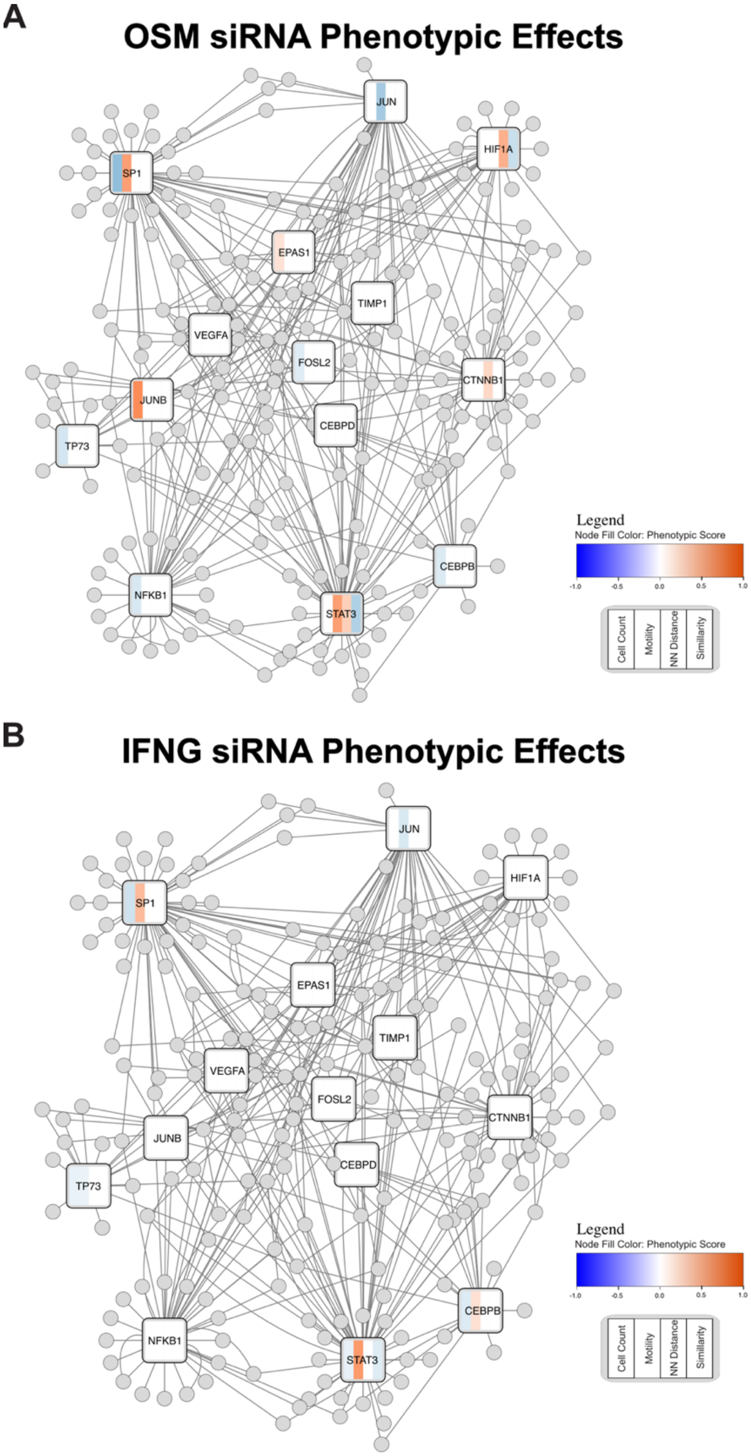


**Supplemental Figure 4: Overview of phenotypic effects of siRNA knockdown**

(A-B) Integrated molecular network of OSM treated cells. Nodes nominated for siRNA screening are labelled by the phenotypic effect of knockdown in the OSM condition (A) and IFNG condition (B) as compared to scramble control. Non-significant changes in phenotype are colored white.

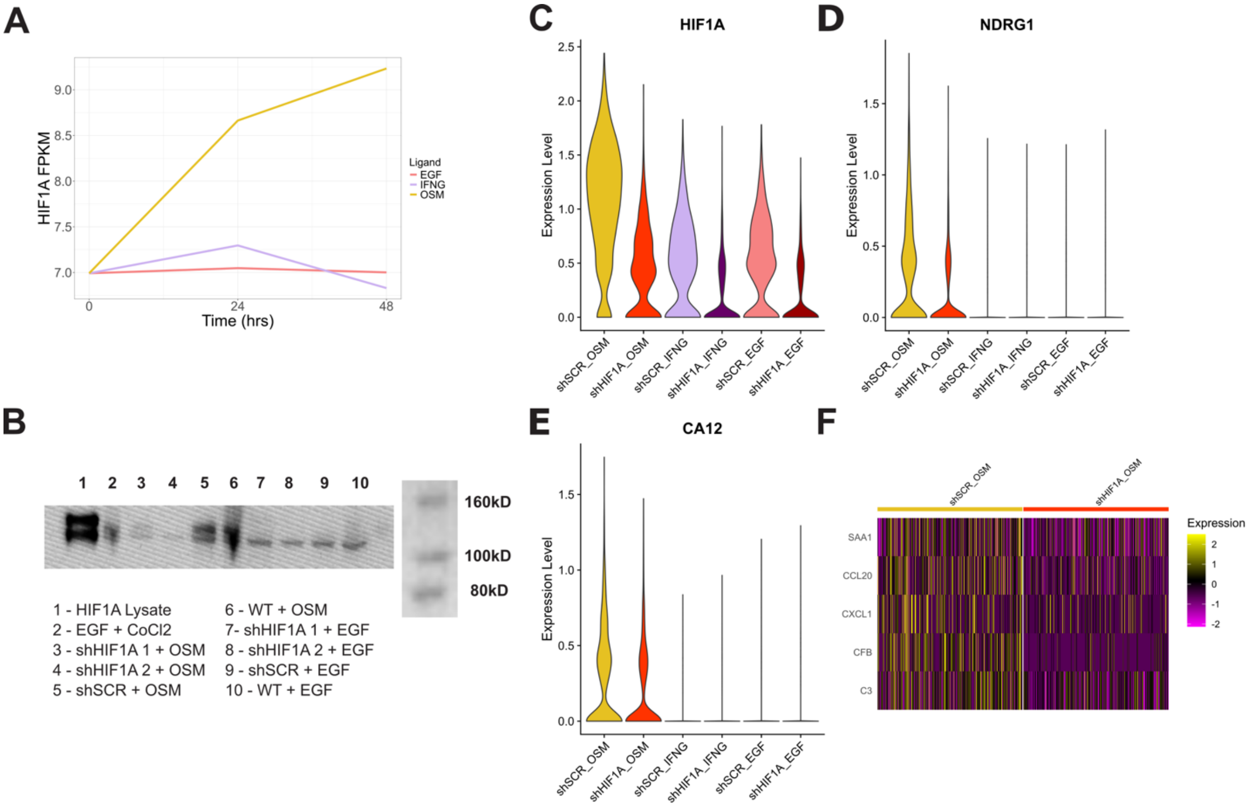


**Supplemental Figure 5: scRNAseq supplemental figures**

(A) Bulk RNAseq demonstrates that OSM uniquely upregulates HIF1A transcriptionally (n=3). (B) Western blot confirming HIF1A knockdown in shHIF1A cell lines. (C) HIF1A expression is significantly reduced in shHIF1A cell lines compared to shSCR. (D-E) Relative gene expression of NDRG1 and CA12: genes that are canonically activated by HIF1A. (E) Genes in the neutrophil chemotaxis Gene Ontology gene set that are upregulated in OSM treated shSCR cells.

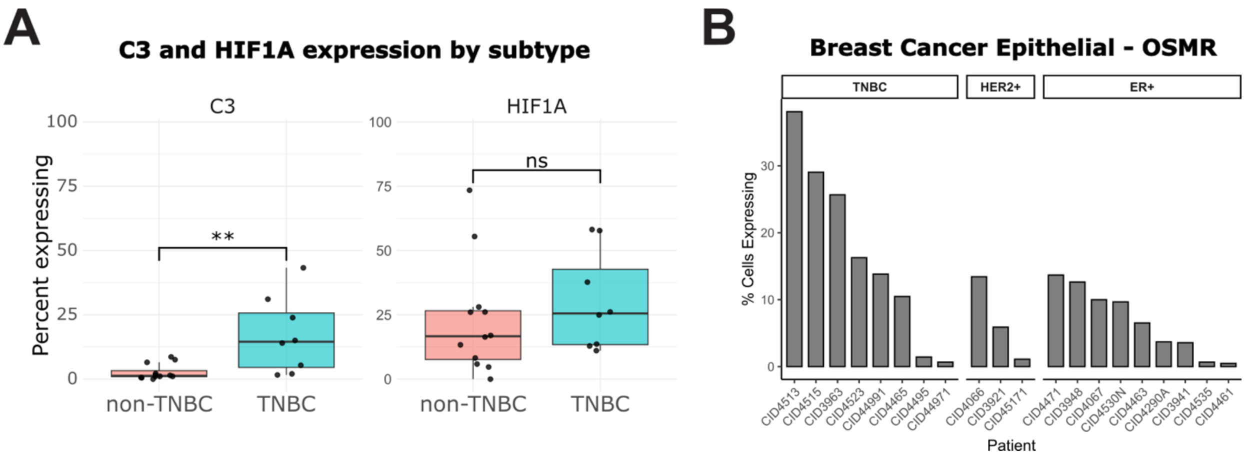


**Supplemental Figure 6: The OSM–HIF1A signaling axis is active in basal-like breast cancer and associated with poor prognosis**

(A) Boxplot showing the percentage of epithelial cells expressing HIF1A and C3 in individual patients. Significance is shown as follows: (* p-value < .05, ** p-value < .01). (B) Barplot showing the percentage of epithelial cells expressing OSMR for each individual patient.
